# GeNETop: Context-Specific Genome-Scale Constrained Models Using Network Topology, Flux Variability, and Transcriptomics

**DOI:** 10.64898/2026.03.16.712013

**Authors:** D. Troitiño-Jordedo, A. Mansouri, R. Minebois, A. Querol, D. Remondini, E. Balsa-Canto

## Abstract

Context-specific genome-scale metabolic models are critical tools for studying cellular metabolism under dynamic conditions. However, most existing methods for deriving these models are designed for steady-state settings and may fail to preserve reactions required for transient metabolic shifts, thereby limiting their compatibility with dynamic FBA.

Here, we present GeNETop, a methodology for deriving context-specific GEMs designed to preserve dynamic compatibility. GeNETop integrates flux variability analysis (FVA), network topology metrics based on the Integrated Value of Influence (IVI), and transcriptomic data to identify reactions that are both flux-flexible and structurally influential. Reactions are prioritized based on variability and maximality indices, while topology and gene expression guide further refinement, reducing dependence on fixed expression thresholds.

Using batch fermentation of *Saccharomyces cerevisiae* as a case study, we evaluate GeNETop against established methods for context-specific metabolic reconstruction.

The resulting networks remain dynamically feasible across growth phases, capture key metabolic transitions, reduce non-essential reactions, and maintain computational tractability.

Overall, GeNETop enables context-specific metabolic reconstructions that are compatible with dynamic simulations while maintaining computational efficiency. By overcoming key limitations of existing approaches, the method supports a more accurate representation of time-dependent metabolic processes in biotechnology and systems biology.

**Author summary:** Cellular metabolism relies on complex networks of reactions to process nutrients, generate energy, and build essential compounds for biomass. Context-specific metabolic models aim to represent only the reactions active under a given condition, improving biological realism and reducing computational complexity in flux balance analysis simulations.

However, metabolic activity adapts dynamically to changing environmental conditions, and reactions that are inactive at one stage may become essential at another.

Many current reconstruction methods are designed for steady-state conditions and may exclude reactions that are required during metabolic transitions, thereby limiting their ability to describe dynamic behavior.

Here, we introduce GeNETop, a novel approach that refines context-specific networks by integrating multiple layers of information. GeNETop identifies the most relevant reactions by considering their flexibility, importance within the network topology, and gene activity levels. In this way, the method generates biologically meaningful models that focus on metabolic pathways relevant under dynamic conditions.

We tested GeNETop on yeast fermentation, a key process in food and biofuel production. The resulting models capture metabolic changes over time and enable stable dynamic simulations, supporting improved flux balance analysis of time-dependent metabolic processes.

## Introduction

Constraint-based genome-scale models (GEMs) have become increasingly popular in biotechnology, providing insights into cellular metabolism under different conditions [1]. These models reconstruct metabolic networks based on genome annotations and apply stoichiometric and mass-balance principles to estimate intracellular flux distributions. GEMs have been widely used to study metabolic processes, including biofuel, bioplastic, pharmaceutical, and food production [2–4].

A widely used approach to predict cellular metabolism with GEMs is flux balance analysis (FBA) [5]. FBA assumes that cells optimize a specific objective function, such as biomass growth, allowing the computation of intracellular flux distributions by solving a constrained optimization problem. However, standard FBA assumes steady-state intracellular metabolism and therefore cannot capture dynamic metabolic shifts. Dynamic FBA (dFBA) extends this framework, integrating a kinetic model to simulate time-dependent metabolic changes [6]. This kinetic model can be formulated as a set of ordinary differential equations that describe the dynamics of crucial extracellular metabolites. This approach provides a comprehensive understanding of the metabolic processes over time [7, 8]. However, even in dFBA implementations, the underlying metabolic network is typically fixed, despite substantial shifts in reaction usage over time.

FBA and dFBA are powerful computational frameworks for simulating cellular metabolism. However, their effectiveness depends on accurate representations of the metabolic network that reflect the actual physiological state of the cell. GEMs, while comprehensive, often include thousands of reactions, many of which are inactive under specific conditions. The large reaction space increases computational complexity and may lead to unrealistic flux distributions and poor interpretability of model predictions. Context-specific metabolic networks (CSNs) address these limitations by tailoring the GEM to a given condition, thereby retaining only the reactions relevant to the condition of interest [9].

Several methods have been developed to tailor genome-scale reconstructions into context-specific networks. One possibility is to integrate additional omics data, such as transcriptomic data. Machado and Herrgård [10] evaluated the performance of the available methods to integrate transcriptomic data into GEMs using published data from *E. coli* and *S. cerevisiae*. The authors concluded that none of the methods outperforms the others in all cases. In any case, GIMME [9] and FASTCORE [11] are probably the most popular methods to reconstruct compact, context-specific metabolic network models.

The GIMME algorithm [9] uses transcriptomic data and required metabolic functionalities to derive CSNs. GIMME removes reactions with expression levels below a user-defined threshold while preserving essential metabolic functions. Its effectiveness in deriving context-specific networks depends on the selected threshold. Recently, Troitiño-Jordedo et al. [12] proposed a compartmentalized GIMME version to allow the definition of different thresholds for user-defined compartments. The added flexibility improved performance.

FASTCORE [11] takes as input the global genome-scale metabolic network and a set of “core” reactions known to be active in the specific context of interest. The algorithm then searches for a flux-consistent subnetwork of the global network that includes all the core reactions and a minimal set of additional reactions. The method is computationally efficient and well-suited for static conditions.

Despite their differences, existing context-specific reconstruction methods share a common assumption: reaction selection is performed under static conditions. In dynamic settings, however, metabolic activity shifts over time, and reactions that appear inactive at one stage may become essential during subsequent phases. Methods that rely primarily on fixed expression thresholds or minimal core sets may therefore exclude reactions required for transient metabolic transitions. As a result, networks derived under steady-state assumptions may not remain compatible with dynamic FBA simulations. Addressing this limitation requires reconstruction strategies that consider not only condition-specific gene expression, but also the potential flux flexibility of reactions and their structural roles within the metabolic network.

These observations highlight the need for reconstruction methods specifically designed to support dynamic metabolic simulations. In this study, we propose GeNETop, a methodology for deriving context-specific genome-scale metabolic networks explicitly designed to preserve dynamic compatibility. GeNETop integrates flux variability analysis (FVA), which identifies reactions capable of operating across a range of fluxes, with network topology measures that capture structural importance, together with transcriptomic data reflecting condition-specific gene activity.

Using yeast batch fermentation as an illustrative example, we demonstrate that integrating these complementary sources of information yields networks that remain feasible across growth phases and better represent metabolic adaptation over time. We compare GeNETop with GIMME and FASTCORE under both steady-state and dynamic scenarios, highlighting differences in reaction retention and dynamic behavior.

## Materials and methods

### Experimental methods for batch fermentation illustrative example

#### Yeast strain

In this study, batch microfermentations were performed using the *Saccharomyces cerevisiae* T73 strain (SC-T73, Lalvin T73, Lallemand, Montreal, Canada).

#### Microfermentations

Microfermentation assays were performed in three independent biological replicates using 500 ml controlled bioreactors (MiniBio, Applikon, The Netherlands) filled with 470 ml of medium (natural grape must). Each bioreactor was inoculated with an overnight starter culture grown in Erlenmeyer flasks containing 25 ml of YPD medium (2% glucose, 0.5% peptone, 0.5% yeast extract) at 25°*C* and 120 rpm in an agitated incubator (Selecta, Barcelona, Spain). Fermentations were inoculated at an initial optical density of OD_600_ = 0.10. Fermentations were monitored by measuring the sugar content by HPLC. Once the sugar content reached 2 g/L, fermentation was complete.

#### Sampling

Extracellular metabolites, including sugars, organic acids, main fermentative by-products, and assimilable yeast nitrogen (YAN) in the form of amino acids and ammonia, were determined at twelve sampling times following the experimental protocol defined in [13]. We also determined the concentrations of higher alcohols and esters for each sampling time. The extraction of volatile compounds and gas chromatography were performed following the protocol of Rojas et al. [14]. Physiological and biomass parameters, including OD600, dry weight (DW), colony-forming units (CFU), and average cell diameter (ACD), were determined at each sampling time, provided that sufficient cell material was available for the corresponding measurements.

#### Transcriptomic analysis

Transcriptomic analysis was performed on cells harvested from the fermentation broth of each biological triplicate at three time points: during the growth phase (T1: 20 hours), at the end of the growth phase (T2: 26.5 hours), and in the early stationary phase (T3: 43.25 hours). To obtain cells, a volume of fermentation broth was collected from the reactor and transferred to a polypropylene tube, which was then centrifuged (4.000 rpm, 5 minutes, 4°*C*) to pellet the cells. The supernatant was discarded, and the tube was flash-frozen in liquid nitrogen and stored at −80°*C* until total RNA was extracted. Following the manufacturer’s protocol, total RNA was extracted using the High Pure RNA Isolation Kit (Roche, Mannheim, Germany). The samples were sequenced using the Illumina Hiseq 2000, paired-end reads 75 bases long, and were deposited under the BioProject ID PRJNA473087. Sequence reads were trimmed and quality filtered using Sickle (minimum read length 50, minimum quality per base 23) and then aligned to the reference genome using Bowtie2 [15]. Gene counts were obtained using HTSeq-count version 0.9.0. [16].

### Theoretical and numerical methods

#### Metabolic reconstruction for the illustrative examples

We used the metabolic reconstruction iMM904 [17] for *S. cerevisiae*, as it was used in [12], to facilitate the model and metabolic analysis in a continuous fermentation process.

For the batch fermentation case, we used the metabolic reconstruction proposed in [2] as an extension of the Yeast8 model [18] (v8.5.0) for *Saccharomyces cerevisiae* T73. This extension integrates reactions pertinent to the synthesis of aromatic compounds, complementing the description of secondary metabolism.

#### Centrality Measures in Network Topology

Centrality measures are used to identify the most important nodes within a network. These measures help quantify the influence or importance of a node based on its position and connections within the network. Centrality measures have been used to analyze networks in a multitude of contexts, from complex transport networks [19] to complex gene networks [20]. There are a number of centrality measures, each designed to capture different aspects of the importance of a node within a network [21]. Common centrality measures include degree centrality (DC), which counts the number of direct connections for each node; betweenness centrality (BC), which measures the extent to which a node lies on the shortest paths between other nodes; ClusterRank (CC, [22]), which quantifies the degree to which nodes in a network tend to cluster together; neighborhood connectivity (NC), which measures how well connected the neighbors of a particular node are within the network; local H index (LHC) which evaluates the influence of a node by considering both the node’s own connections and the connections of its neighbors and collective influence (CIC) [23] which refers to the impact that a group of nodes can have on the overall network, particularly in terms of spreading information or influence. However, relying on a single centrality measure can lead to a misestimation of node importance. For instance, a node with low degree centrality might still be crucial for network connectivity if it has high betweenness centrality.

Therefore, we have used the integrated value of influence (IVI score) [24], which simultaneously considers global, semi-local, and local CMs, making it a suitable metric for capturing all topological characteristics of the reaction network.

The IVI formula combines the most important local (i.e., degree centrality and ClusterRank), semi-local (i.e., neighborhood connectivity and local H-index), and global (i.e., betweenness centrality and collective influence) centrality measures in such a way that both synergize their effects and remove their biases [25]. The IVI score of each node *i* in a network is formulated as follows:

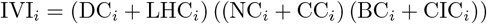

with: *DC*(*v*_*i*_) = N(*v*_*i*_), where N(*v*_*i*_) is the number of direct neighbors of node *v*_*i*_.

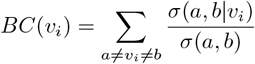

where *σ*(*a, b*) is the number of shortest paths from *a* to*b*, and *σ*(*a, b*|*v*_*i*_) is the number of those paths passing through *v*_*i*_.

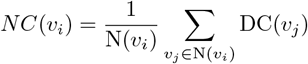

where *v*_*j*_ is a node from the direct neighborhood of *v*_*I*_

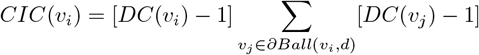

*Ball*(*v*_*i*_, *d*) is the set of nodes inside a ball of radius *d* (shortest path) around *v*_*i*_, and

*∂Ball*(*v*_*i*_, *d*) is the frontier of the ball

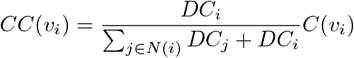

where *C*(*v*_*i*_) is the clustering coefficient of node *v*_*i*_:

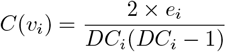

where *e*_*i*_ is the number of edges between the neighbors of node *v*_*i*_.

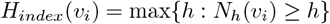

with *N*_*h*_(*v*_*i*_) is the number of neighbors of *v*_*i*_ with a degree ≥ *h*.

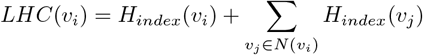

#### Flux balance analysis (FBA)

Flux balance analysis is a constraint-based optimization framework widely used to predict intracellular metabolic fluxes under steady-state conditions [5]. The method relies on a genome-scale metabolic model (GEM), that represents metabolic reactions as a stoichiometric network, in which metabolites are connected through enzymatic reactions. FBA is based on the fundamental assumption that intracellular metabolite concentrations remain constant over time, leading to the steady-state mass balance equation:

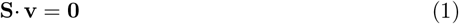

where **S** is the stoichiometric matrix (*m* × *n*), with *m* metabolites and *n* reactions. Each column represents a reaction, and each row represents a metabolite. **v** is the vector of metabolic fluxes (*n*×1), which represent the rates of biochemical reactions.

To ensure feasibility, FBA imposes flux constraints based on thermodynamic, enzymatic, and physiological considerations:

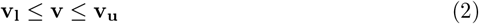

where **v**_**l**_ and **v**_**u**_ define the lower and upper flux bounds for each reaction. Typically, reversible reactions have **v**_**l**_ *<* **0** and **v**_**u**_ *>* **0**, while irreversible reactions have **v**_**l**_ = **0**.

FBA assumes that cells optimize a given objective function *J*(**v**), often chosen to represent a biologically relevant process such as biomass growth. The problem is then formulated as an optimization problem as follows:

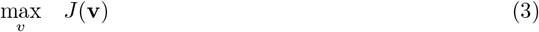

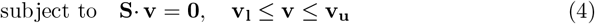

Note that in many cases, multiple flux distributions can achieve the same optimal objective function value, leading to degeneracy in the solution space. To address this, parsimonious FBA (pFBA) introduces an additional optimization step that minimizes the total sum of fluxes, assuming that cells utilize the least enzymatic resources necessary to achieve a given objective [26]. Minimizing the total flux yields solutions that favor a lower enzymatic burden while still achieving optimal growth or production objectives.

#### Flux variability analysis (FVA)

Flux Variability Analysis extends FBA by exploring the range of possible flux values for each reaction while maintaining a given level of the objective function [27]. Unlike standard FBA, which provides a single optimal flux distribution, FVA identifies alternative feasible solutions, revealing the flexibility of the metabolic network.

This is achieved by solving two linear programming (LP) problems for each reaction:

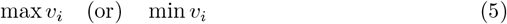

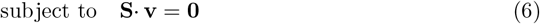

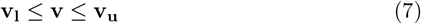

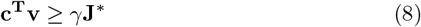

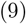

where *v*_*i*_ is the flux of interest, *J*^*∗*^ is the optimal value of the objective function as obtained in the FBA, *γ* is a user-defined threshold that ensures that the system operates near the optimal solution.

By independently maximizing and minimizing each flux *v*_*i*_, FVA determines the minimum and maximum feasible values of each flux, thus identifying reactions with high or low flexibility. If flux variability is zero, the reaction operates at a fixed rate and is therefore considered essential; if flux variability is nonzero, alternative metabolic pathways may compensate for the reaction; and if both minimum and maximum values are zero, the reaction is inactive.

#### Dynamic flux balance analysis (dFBA)

Dynamic Flux Balance Analysis (dFBA) extends standard FBA by incorporating time-dependent changes in extracellular metabolite concentrations. Unlike traditional FBA, which assumes steady-state intracellular fluxes, dFBA accounts for the temporal evolution of metabolic processes, making it well suited for modeling dynamic biological systems such as batch or continuous fermentations [6]. dFBA combines the stoichiometric constraints of FBA with a kinetic model that describes the dynamics of extracellular metabolite concentrations. This kinetic model can be formulated as a set of ordinary differential equations (ODEs):

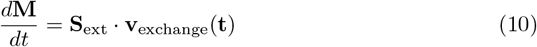

*v*_exchange_(*t*) represents the exchange fluxes for extracellular metabolites, *S*_ext_ is the stoichiometric matrix for extracellular metabolites, and **M** is the vector of extracellular metabolite concentrations.

Since dFBA introduces time-dependent constraints, specialized numerical approaches are required to couple FBA with the kinetic equations for extracellular metabolites.

Note that in the direct approach, an FBA problem is solved at each time step during the numerical solution of the kinetic model, using reliable implicit ODE integrators with adaptive step size and error control. The numerical solution reads as follows:

At each *t*_*j*_ time the corresponding FBA problem is solved, that is:

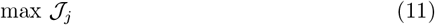

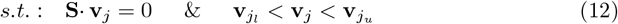

with subindex *j* referring to time *t*_*j*_.

Note that the constraints for external fluxes are defined taking into account the dynamic model Eq. (10) as follows:

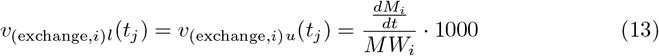

where *MW*_*i*_ denotes the molecular weight of the corresponding compound *i*.

#### Dynamic model for the batch fermentation illustrative example

A dynamic kinetic model was used that is suitable for explaining nitrogen-limited batch fermentations. The model was originally introduced by Moimenta et al. [8], and its formulation is only briefly summarized here.

Biomass (*X*) is modeled based on its major components: carbohydrates (*X*_*C*_), protein (*X*_*P*_), and mRNA (*X*_*mRNA*_), with their dynamics given by:

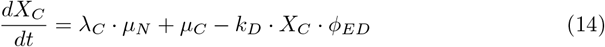

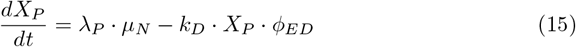

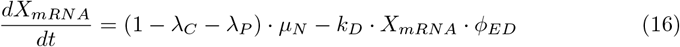

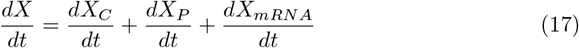

where *λ*_*C*_ = 0.29, *λ*_*P*_ = 0.59, and 1 −*λ*_*C*_ − *λ*_*P*_ = 0.12 are the biomass composition fractions [28]. The model differentiates between primary growth, which occurs in the presence of nitrogen (with a growth rate *µ*_*N*_), and secondary growth (with a growth rate *µ*_*C*_), attributed to the accumulation of carbohydrates following nitrogen depletion [29]. The term *k*_*D*_ *ϕ*_*ED*_ represents the decay induced by ethanol.

Yeasts use nitrogen and sugars to grow, with assimilable nitrogen (amino acids and ammonium) as the limiting substrate. A generalized mass action model describes the transport of nitrogen sources:

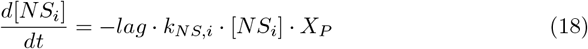

where we denote [*NS*_*i*_] the concentration of each nitrogen source, that is, amino acids and ammonia, and 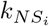the associated kinetic parameter. The lag model follows the Baranyi-Roberts model [30].

Glucose and fructose transport is modeled using Michaelis-Menten kinetics, with their uptake decreasing during nitrogen starvation.

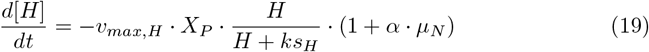

where *v*_*max,H*_ corresponds to the maximum uptake rate for the given hexose (glucose or fructose), *ks*_*H*_ corresponds to the Michaelis-Menten constant. The term *α · µ*_*N*_ describes the increase in sugar transporter production during the exponential phase.

The transition to the stationary phase is modeled as:

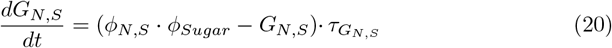

where *ϕ*_*N,S*_ and *ϕ*_*Sugar*_ are sigmoid functions that control responses to nitrogen and sugar depletion.

During fermentation, glucose and fructose are converted into several metabolites.

Their excretion rate is directly proportional to glucose and fructose uptake:

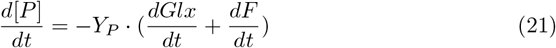

The yield coefficient (*Y*_*P*_, where*P* represents the given metabolite expressed in (g_i_/g_sugar_) varies significantly depending on the fermentation phase. Note that production models vary by release phase: proportional to sugar uptake (ethanol), gradually regulated (glycerol, acetate, lactate), delayed until nitrogen is depleted (succinate, ethyl acetate, isoamyl acetate, isobutanol), or delayed and slowly repressed during the stationary phase (ethyl acetate, phenyl acetate, 2-phenyl ethanol, 2-3 butanediol). We use the smooth functions *ϕ*_*N,S*_ and *G*_*N,S*_ to accommodate the different scenarios. Detailed equations can be found in [8].

#### Software tools

Network topology analyses and the computation of centrality measures were performed using the R implementation described in [24].

Constraint-based metabolic analyses were carried out using the COBRA Toolbox [31] in MATLAB. Flux balance analysis (FBA), parsimonious FBA (pFBA), flux variability analysis (FVA), and the context-specific reconstruction algorithms GIMME [9] and FASTCORE [11] were implemented using this framework. Linear programming problems were solved using the IBM ILOG CPLEX solver (version 22.1.0.0) [32].

The dynamic kinetic model used in the batch fermentation case study was calibrated using the AMIGO2 toolbox [33]. This toolbox was also used to implement the direct approach for solving the dynamic flux balance analysis (dFBA) problem by coupling the kinetic model with the constraint-based optimization framework.

## Results

### GeNETop workflow

GeNETop requires a genome-scale metabolic reconstruction together with information constraining metabolism under dynamic conditions, including time-resolved extracellular measurements or an equivalent kinetic description of biomass and exometabolome dynamics, as well as transcriptomic measurements collected at representative sampling times. The workflow integrates flux variability analysis (FVA), network topology through the integrated value of influence (IVI), and transcriptomic data. Figure 1A summarizes the overall procedure.

**Fig 1.**
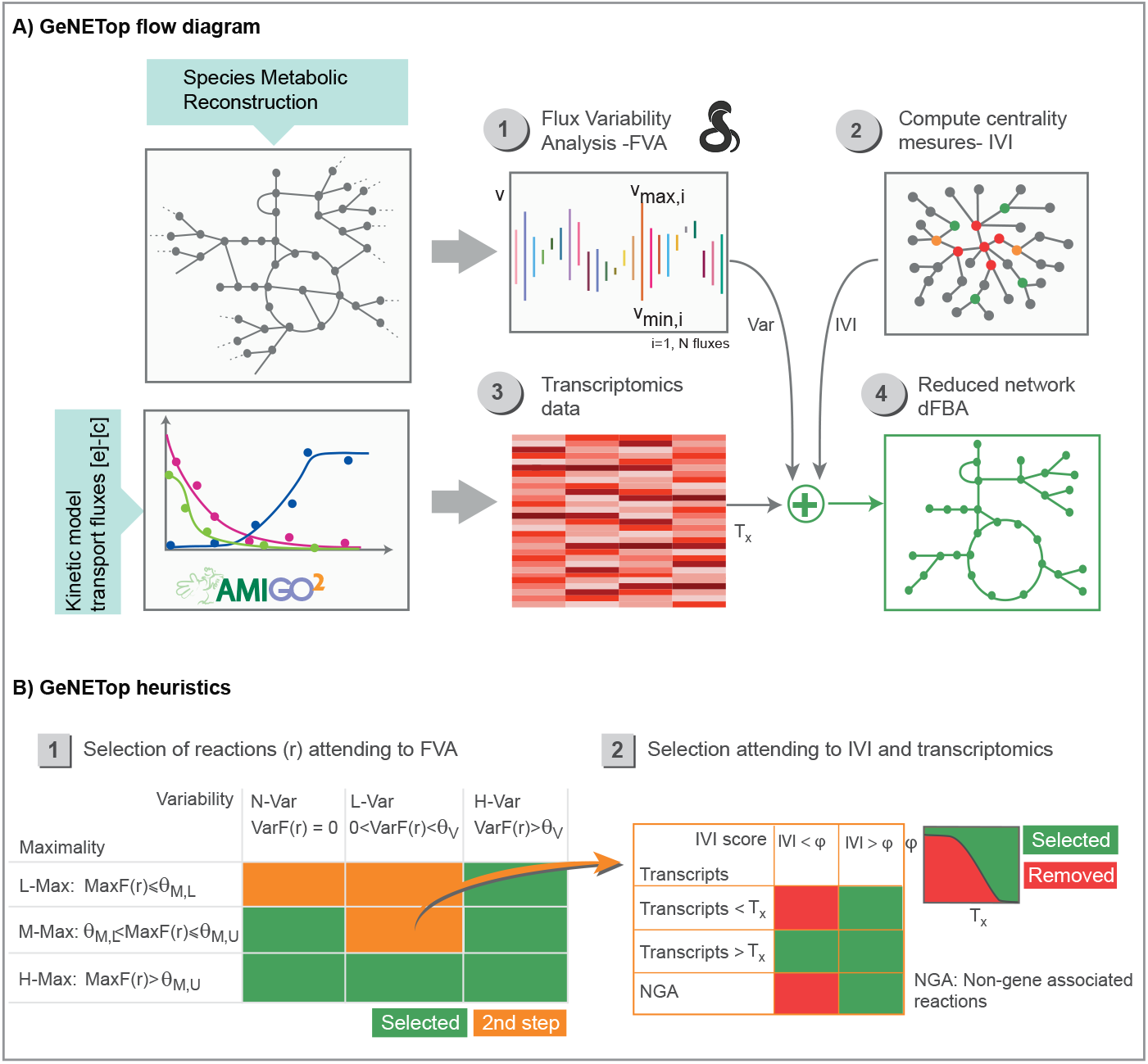
GeNETop flow diagram and heuristics. Figure A) presents the elements required by GeNETop and their interactions. Figure B) illustrates the heuristics used to identify relevant context-specific reactions using flux variability analysis, the integrated value of influence (IVI score), and transcriptomics data.

The method proceeds in four steps: (1) classification of reactions according to dynamic flux variability analysis (FVA); (2) ranking reactions according to their integrated value of influence (IVI); (3) formulation of a reduced context-specific network by integrating IVI rankings with transcriptomic data; and (4) solution of the dFBA problem using the reduced model. Steps 1 and 2 are independent and can therefore be performed in parallel. In the sequel, we briefly describe the four steps; interested readers can find further details in the Supplementary File Section 1.

### Step 1: Classifying metabolic reactions using FVA

In the first step, reactions are classified according to their flux flexibility under the dynamic constraints of the system. External transport fluxes are constrained using the dynamic model, and FVA is performed at the sampling times for which transcriptomic data are available.

For each reaction *r*, the maximum and minimum feasible fluxes (*v*_*max,r*_ and *v*_*min,r*_) are obtained and used to compute two metrics: the *variability index, V arF* (*r*) = |*v*_*max,r*_ − *v*_*min,r*_|, representing the allowable flux range, and the *maximality index, MaxF* (*r*) =| *v*_*max,r*_| + |*v*_*min,r*_|, representing the maximum potential flux magnitude.

Reactions are classified according to the variability and maximality indices as follows:

- *Non–variable reactions (N–Var)*: Reactions whose variability index is zero, that is, *V ar*(*r*) = 0.
- *Low variable reactions (L–Var)*: Reactions whose variability index is low (0 *< V ar*(*r*) *< θ*_*V*_).
- *High variability reactions (H–Var)*: all remaining reactions.
- *Low maximality reactions (L–Max)*: Reactions whose maximality index is below a given threshold, *MaxF* (*r*) ≤ *θ*_*M,L*_.
- *Medium maximality reactions (M–Max)*: Reactions whose maximality index is within the range *θ*_*M,L*_ *< MaxF* (*r*) ≤ *θ*_*M,U*_.
- *High maximality reactions (H–Max)*: all remaining reactions.

The threshold parameters *θ*_*V*_, *θ*_*M,L*_, and *θ*_*M,U*_ are chosen to distinguish numerical artefacts from biologically meaningful flux values. The variability threshold *θ*_*V*_ corresponds to the smallest flux variation distinguishable from solver numerical tolerance, while the maximality thresholds define lower and upper ranges separating negligible fluxes from potentially active reactions. Default values (*θ*_*V*_ = 10^−10^, *θ*_*M,L*_ = 10^−10^, *θ*_*M,U*_ = 10^−7^) were selected to ensure robustness while avoiding misclassification of numerical artefacts.

As illustrated in Figure 1B, reactions exhibiting high variability are automatically retained in the context-specific network regardless of maximality. Among non-variable reactions, those with medium or high maximality are also retained. Within the low-variability group, only reactions with high maximality are automatically selected. Remaining reactions are further evaluated in the next step using network topology and transcriptomic information.

### Step 2: Ranking metabolic reactions using IVI

In the second step, reactions are ranked according to their topological importance within the metabolic network. A bipartite network linking reactions and metabolites is first constructed, then transformed into a reaction network in which nodes correspond to reactions and edges connect reactions that share at least one metabolite.

To preserve meaningful topological structure, highly connected currency metabolites (e.g., ATP, NAD(H), water) are removed, preventing the creation of artificial connections between otherwise unrelated reactions.

The integrated value of influence (IVI) is then calculated for each reaction. IVI combines several centrality measures, including Degree, Betweenness, Neighborhood Connectivity, Collective Influence, Local H-Index, and ClusterRank, into a single metric capturing multiple dimensions of node importance. This integration provides a robust ranking of reactions within the metabolic network.

### Step 3: Integrating IVI and transcriptomics to reduce the network

The third step constructs the context-specific network by combining IVI rankings with transcriptomic data for reactions that were not selected in Step 1 (orange reactions in Figure 1B).

Reactions with IVI values below a threshold *φ* are removed unless their transcription level exceeds a threshold *T*_*X*_, defined as the *X*th percentile of transcript abundance. This rule ensures that reactions with low topological influence are retained when strongly expressed in the given context.

The threshold *φ* depends on both reaction variability and IVI values and is defined as follows:

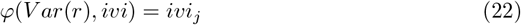

where *ivi* is the ordered set of IVI values and

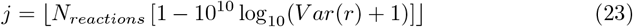

with *N*_*reactions*_ the total number of reactions in the original GEM.

The threshold *φ* dynamically links reaction variability to network topology, imposing stricter IVI requirements for reactions with low flux variability while relaxing the criteria for reactions with greater flexibility. This formulation creates a dynamic inclusion threshold that balances network topology and gene expression. Importantly, the approach naturally retains non-gene-associated reactions when their IVI values indicate structural relevance. The rationale behind this heuristic is discussed in the Supplementary File.

### Step 4: Dynamic flux balance analysis with the reduced network

The final step solves the dFBA problem using the reduced context-specific network. The kinetic model describing extracellular dynamics is given by

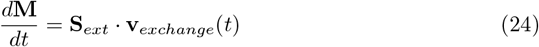

where **M** represents extracellular metabolite concentrations and **S**_*ext*_ the stoichiometric matrix for extracellular metabolites.

At each time point *t*_*j*_, an FBA problem is solved using the reduced stoichiometric matrix **S**_**r**_:

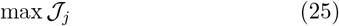

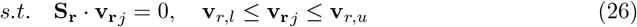

Although GeNETop is designed for dynamic environments, we first verified its compatibility with previously reported steady-state reconstruction approaches using the continuous fermentation dataset described in [12]. Structural comparisons with the published cGIMME configuration under identical constraints and transcriptomic data are provided in Supplementary File Section 2 and Supplementary Table S1. Consistent steady-state flux predictions confirmed that GeNETop remains compatible with established context-specific modeling frameworks.

Additional methodological details and an extended rationale for each step of the GeNETop workflow are provided in Supplementary Section S1.

### Illustrative batch fermentation case study in *S. cerevisiae* T73

To illustrate the performance of GeNETop, we considered a representative batch yeast fermentation experiment using the *Saccharomyces cerevisiae* T73 strain. Biomass, carbon, and nitrogen sources, and metabolic products, including ethanol, glycerol, succinate, several carboxylic acids, and aromatic compounds, were measured at approximately twelve sampling times throughout the process. Transcriptomic data were collected at three representative time points: 20 *h*, 26.5 *h*, and 43.25 *h*, corresponding to the exponential phase, the growth-no growth transition, and the stationary phase, respectively.

### GeNETop reconstruction of the T73 context-specific network

The Yeast8 metabolic reconstruction [18] was used as reference, comprising 4060 reactions. External fluxes were obtained by calibrating the kinetic model proposed by [8] against biomass and exometabolomic time-series data (see Materials and Methods). The calibrated model describes biomass growth, uptake of sugars, ammonium, and amino acids, and production of ethanol, glycerol, succinate, citrate, malate, lactate, isobutanol, isoamyl alcohol, 2-phenylethanol, acetyl, isobutyl acetate, isoamyl acetate, and phenylethyl acetate. The corresponding fits are shown in the Supplementary File Section 3.

The first step of GeNETop consists of flux variability analysis. Using biomass growth as the cellular objective, variability and maximality values were computed for each reaction at the transcriptomic sampling times (Supplementary Table S2.1), and reactions were classified into the categories N-Var, L-Var, H-Var, L-Max, M-Max, and H-Max for subsequent selection. This step automatically retained 2381 reactions, while the remaining ones were evaluated in the following steps.

The second step ranks reactions according to their integrated value of influence. IVI calculation starts from a bipartite metabolic network linking reactions and metabolites, from which a reaction network is derived by connecting reactions that share at least one metabolite. To avoid overconnecting the network through functionally uninformative hubs, prevalent metabolites such as ATP, ADP, NAD, NADH, NADP, NADPH, *O*_2_, *H*_2_*O*, PI, *H*^+^, *Glu*_*L*_, and *CO*_2_ were excluded. Centrality measures and IVI were then computed for each reaction (Supplementary Tables S2.2 and S2.3). The IVI distribution decreases rapidly: after excluding highly connected hubs and reactions embedded in long chains, most reactions (99.83%) show IVI values between 1 and 5. The highest-ranking reactions are biologically meaningful.

The most influential reaction is the hydrolysis of diphosphate to phosphate (r 0568, IVI=100), a central process linked to ATP and nucleic acid metabolism and, therefore, to energy transfer and redox balance. The second-ranked reaction, r 0112 (IVI=80.1), catalyzes the conversion of acetate, ATP, and coenzyme A into acetyl-CoA, AMP, and diphosphate in the cytoplasm. This reaction links acetate utilization to the synthesis of lipid, sterol, and biomass precursors and is particularly relevant during late fermentation stages, when glucose becomes scarce, and acetate or ethanol accumulate. The third-ranked reaction, r 2194 (IVI=47.5), is involved in fatty acyl-CoA synthesis and is therefore directly associated with lipid metabolism and membrane biosynthesis. Subsequent high-IVI reactions are involved in the biosynthesis of cellular components, including glycoproteins, lipids, and amino acids, which are essential for growth and survival.

In the third step, IVI and transcriptomic information are combined to determine the selection of reactions that were not retained in the first step through the *φ* function defined in Eq. (22). Transcriptomic data are reported in Supplementary Table S2.4 and the resulting selection criteria in Table S2.5. This step retained an additional 363 reactions, yielding a final GeNETop context-specific network with 3018 reactions. The complete selection procedure required approximately 12 minutes on a standard desktop computer (Intel® Core− i7-9750H CPU @ 2.60 GHz × 12), with most of the computational cost associated with FVA, which requires solving two optimization problems at each of the three sampling times.

To assess the robustness of reaction selection with respect to the threshold parameters, we performed a sensitivity analysis by varying the flux-variability threshold to the values 10^−10^, 10^−7^, and 10^−5^. The resulting networks showed only moderate variations in size and composition (below 2% of the total number of reactions), indicating that the overall CSN structure is relatively stable with respect to this parameter. However, the most restrictive threshold (10^−5^) led to convergence issues in dFBA, suggesting that limited transcriptomic temporal resolution may hinder the retention of reactions required for intermediate metabolic transitions. A detailed summary of this analysis is provided in Supplementary Table S2.5 and the Supplementary File Section 4.

### Comparison with GIMME and FASTCORE under static conditions

We next assessed the performance of GeNETop relative to two widely used algorithms, GIMME [9] and FASTCORE [11]. These methods pursue different objectives: GIMME emphasizes agreement with gene expression data and predefined metabolic functions, FASTCORE prioritizes speed and compactness, whereas GeNETop was specifically formulated for dynamic settings. To compare the three approaches under identical conditions, we considered the exponential phase at *t* = 21 *h*, where biomass maximization is a suitable FBA objective.

GIMME was supplied with the same information as GeNETop, namely the 38 externally constrained fluxes obtained from the kinetic model and transcriptomic data for all gene-associated reactions at *t* = 21 *h*. Using a threshold of 1000 transcripts, corresponding to the 65th percentile and 908 reactions, GIMME produced a CSN with 2878 reactions. Notably, GIMME incorporated all reactions lacking transcriptomic information, regardless of whether they were gene-associated, adding 1438 reactions. These included many transport reactions, lipid metabolism, complex lipid degradation into fatty acids, and membrane biosynthesis pathways. In addition, to ensure convergence of the FBA problem, GIMME incorporated 118 additional reactions with transcript values ranging from 31 to 996. The selected reactions are listed in Supplementary Table S2.6.

FASTCORE was first used in a black-box mode by specifying the reactions corresponding to measured extracellular metabolites (FASTCORE-E), so that the algorithm would return the minimum set of reactions compatible with the available exometabolomic data. This produced a CSN containing 390 reactions in approximately 15 seconds. We also tested a second configuration, FASTCORE-T, in which reactions with transcriptomic support were explicitly included. This resulted in a larger CSN with 676 reactions, although the algorithm reported the network as inconsistent.

GeNETop and GIMME produced substantially larger context-specific networks than FASTCORE-E and FASTCORE-T. GeNETop generated the largest CSN, corresponding to a 26% reduction relative to the original Yeast8 reconstruction, whereas GIMME achieved a 44% reduction. FASTCORE-E produced a highly compact network, but at the cost of excluding numerous transcript-supported reactions, while FASTCORE-T yielded a somewhat larger yet inconsistent reconstruction.

### Dynamic comparison of GeNETop and GIMME

We further compared GeNETop and GIMME under dynamic conditions, using the kinetic model describing biomass, substrate, and product dynamics to constrain the dFBA problem.

GeNETop was designed to identify a context-specific network that retains reactions potentially required across different fermentation phases. The dFBA implementation employed a time-varying objective function that balances energy generation and protein turnover after nitrogen depletion [8]. Under these conditions, the GeNETop-derived CSN yielded successful dynamic simulations. The corresponding flux scores, computed as time integrals of the dynamic fluxes, are reported in Supplementary Table S3.1.

To apply GIMME under the same conditions, we merged the CSNs reconstructed independently from transcriptomic data across different sampling times into a single expanded network. Because transcript levels vary substantially over time, a fixed threshold of 1000 transcripts would be overly restrictive, especially during the stationary phase. For this reason, we instead used the 50% percentile at each phase, corresponding to approximately 800 transcripts in the exponential phase, 500 during the growth-no growth transition, and 400 in the stationary phase. The resulting merged CSN was then used in dFBA. Under these conditions, GIMME showed convergence problems, and simulations could only be run between 17 *h* and 79.5 *h*. The corresponding flux scores are reported in Supplementary Table S3.2.

To investigate the origin of these discrepancies, we compared the flux distributions predicted by GeNETop and GIMME at 17 *h* and 79.5 *h* (Supplementary Tables S3.3 and S3.4). Figure 2A shows the overlap and method-specific reactions at both time points. At 17 *h*, GeNETop used 489 reactions, and GIMME used 510, of which 460 were shared, indicating substantial agreement in the early fermentation phase. At 79.5 *h*, GeNETop predicted 98 reactions with non-zero flux, and GIMME predicted 96, with 82 reactions shared. For both methods, the smaller number of active reactions at 79.5 *h* is consistent with lower metabolic activity during the stationary phase.

**Fig 2.**
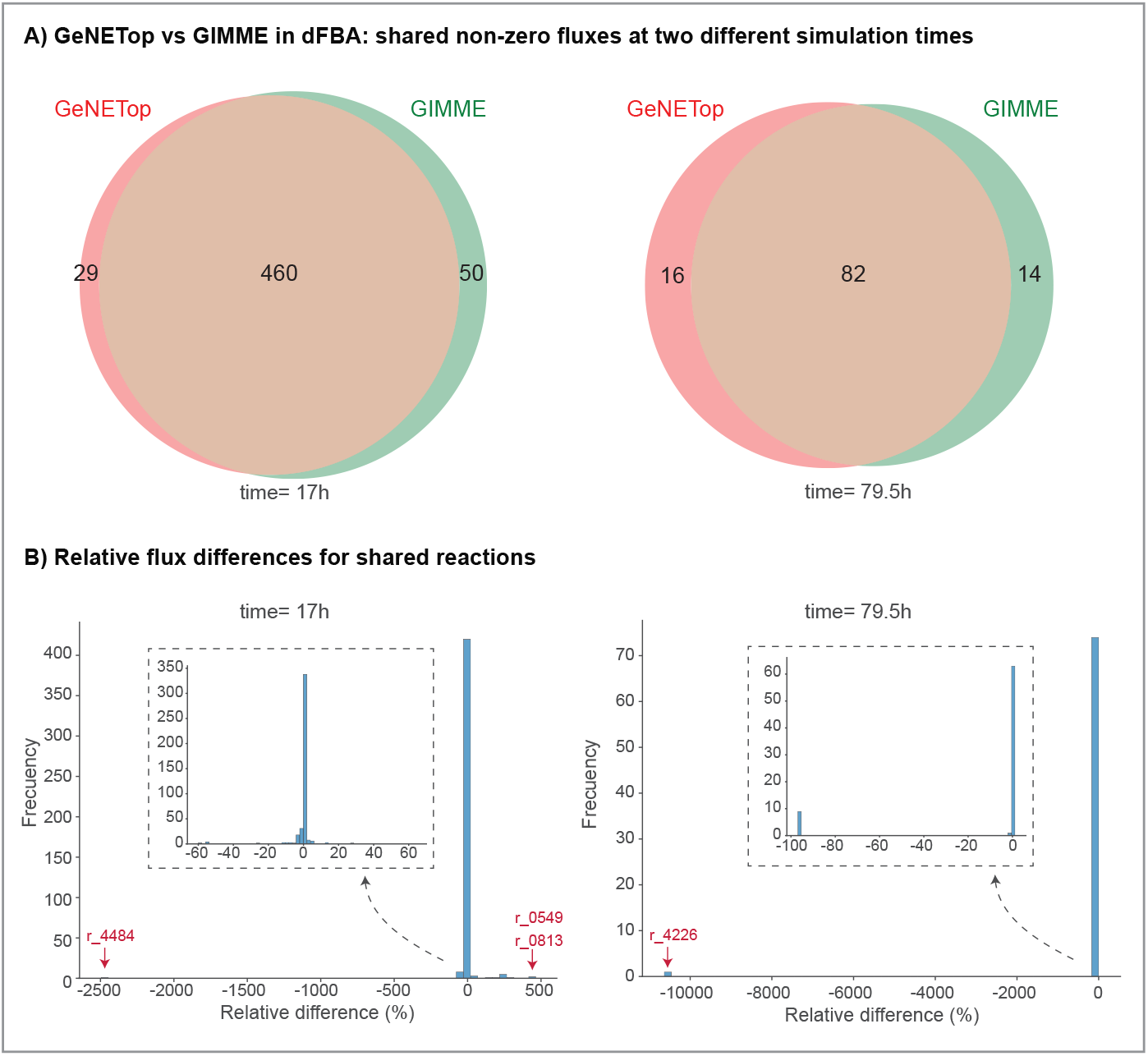
GeNETop vs GIMME in dynamic flux balance analysis. A) The Venn diagrams illustrate the overlap and unique reactions used by GeNETop and GIMME at t=17 h and t=79.5 h. The figures highlight shared core reactions (overlap) and method-specific reactions, reflecting the differences in algorithmic approaches. GIMME uses more reactions than GeNETop at t=17 h, but the number of reactions is practically identical at t=79.5 h. The number of reactions decreases over time, indicating reduced metabolic activity. B) The histograms represent the relative flux differences (in %) for shared reactions between GeNETop and GIMME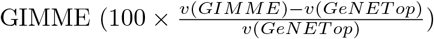. These plots show that most shared reactions exhibit small relative differences in flux predictions between GeNETop and GIMME, clustering around 0%. This indicates a strong overall agreement between the two methods for most reactions.

The distribution of relative flux differences for shared reactions is shown in Figure 2B. At both time points, most shared reactions cluster around 0%, indicating broad agreement between the methods. However, the proportion of large deviations increases at 79.5 *h*, revealing greater divergence in predictions during the stationary phase.

A pathway-level analysis of the fluxes by compartment and metabolic function (Supplementary Tables S3.3 and S3.4) revealed several notable differences. One example concerns pyruvate routing: GeNETop predicts pyruvate transport into mitochondria through r 2034 (flux = 0.376), whereas GIMME predicts flux through the D-lactate/pyruvate antiport r 1138 (flux = 0.371). Reaction r 2034 was excluded from the GIMME CSN because its transcript abundance (436) was well below the threshold of 1000, while r 1138, which lacked transcript information, was incorporated to ensure network completeness. Since pyruvate links cytosolic glycolysis to the mitochondrial tricarboxylic acid cycle, this difference has direct consequences for compartmentalized metabolic interpretation.

A second difference concerns the mitochondrial reduction of acetaldehyde to ethanol in GeNETop through reaction r 0165 (flux = 0.245), a process coupled to NADH oxidation and therefore relevant for redox balancing. GIMME did not predict this flux, despite the high transcript abundance of the associated reaction across the three sampling times (17292, 4358, and 1891), because the required transport reactions r 1763 and r 1632 lacked transcript information and were incorporated only through network completion. In contrast, GeNETop retained all three reactions directly after the FVA-based filtering step.

Differences were also observed in the hydration of carbon dioxide to bicarbonate. GeNETop predicts this process directly in the cytosol through r 1667 (flux = 0.884), whereas GIMME routes it through a more complex sequence involving the transport of CO_2_ to the nucleus (r 1694), hydration in the nucleus (r 1665), and the transport of bicarbonate back to the cytosol (r 1669). Reaction r 1667 was selected by GeNETop after FVA but excluded by GIMME because of its transcript abundance (993, 597, and 266 across the three time points), whereas the alternative reactions in GIMME were introduced through completion rules because transcript information was unavailable.

Another clear distinction involves reactions r 4484 and r 4274. In GIMME, r 4484 consumes succinate to produce O-succinyl-L-homoserine, which is subsequently hydrolyzed by r 4274 to regenerate succinate. Taken together, these reactions form a loop that appears to contribute to 2-oxobutanoate production. GeNETop does not use r 4274; instead, it predicts 2-oxobutanoate production through the deamination of L-threonine via r 0692, which was excluded from GIMME because its transcript abundance never exceeded 65. Reactions r 4484 and r 4274 were retained in GIMME because their transcript abundance during the exponential phase was 1898, while all three reactions were retained by GeNETop after the FVA-based selection step.

At 79.5 *h*, when overall metabolic activity is markedly reduced, differences in CO_2_ hydration persist, although at much lower flux values. Most of the remaining discrepancies arise from transport reactions between compartments and the reactions occurring within those compartments. For example, GIMME predicts the transport of L-alanine and L-glycine from the cytosol to the vacuole, where they are combined into the Ala-Gly dipeptide and water through r 4203, followed by the transport of the dipeptide back to the cytoplasm and release into the medium. GeNETop does not predict flux through these reactions at this time point. Nevertheless, both methods predicted the use of these reactions during exponential growth, with differences of less than 0.6%.

## Discussion

This work introduces GeNETop, a methodology for deriving context-specific metabolic networks tailored to dynamic environments. By integrating flux variability analysis (FVA), network topology metrics (via IVI scores), and transcriptomics, GeNETop addresses the challenge of deriving context-specific networks that remain biologically consistent under changing environmental conditions. The method enables compatibility with dynamic flux balance analysis (dFBA) while reducing dependence on dense time-series transcriptomic datasets and maintaining computational efficiency. Unlike previous context-specific reconstruction approaches that rely primarily on transcriptomic thresholds or predefined core reaction sets, GeNETop explicitly incorporates flux flexibility and network topology to preserve reactions that may become relevant during metabolic transitions.

To evaluate its behaviour under steady-state conditions, we first compared GeNETop with the previously published cGIMME approach using the continuous fermentation dataset reported in that work [12]. Although the two methods generated reduced networks with noticeable structural differences, parsimonious FBA produced identical optimal flux predictions for both reconstructions. This result indicates that GeNETop remains consistent with established context-specific reconstruction approaches under steady-state conditions and provides a framework for supporting dynamic simulations.

To assess its performance in batch fermentation processes, we applied GeNETop and compared its outputs with those obtained using GIMME [9] and FASTCORE [11].

Under static conditions, both GeNETop and GIMME produced feasible context-specific networks. However, the resulting CSNs differed structurally, reflecting the distinct principles underlying the algorithms. FASTCORE-E (enforcing external flux constraints) generated an overly minimal CSN, while FASTCORE-T (incorporating transcriptomic information) resulted in an inconsistent network in the present case study. These results illustrate that FASTCORE-based reconstructions may require careful definition of the core reaction set to ensure biological feasibility. This behaviour reflects the strong dependence of FASTCORE on the definition of the core reaction set. When the core set is limited to reactions associated with measured metabolites or transcriptomic evidence, auxiliary reactions required to maintain metabolic connectivity or support dynamic feasibility may be excluded from the reconstruction.

When applying GIMME to dynamic conditions, an additional difficulty arises because transcript levels fluctuate over time, while the algorithm relies on a single threshold applied across all sampling points. Consequently, reactions that appear inactive at one stage but become relevant later in the process may be excluded from the reconstructed network. Depending on the selected threshold, this approach may either retain irrelevant pathways, increasing computational complexity, or remove reactions required to capture metabolic transitions. We explored the alternative strategy of generating independent CSNs for each sampling time and merging them into a unified network. However, this approach is limited by the temporal resolution of transcriptomic sampling and may fail to retain intermediate reactions required for smooth metabolic transitions. Moreover, because GIMME primarily evaluates gene-associated reactions, it may neglect non-gene-associated reactions that still play important roles in metabolic pathways, particularly during transient metabolic states.

Our direct comparison of GeNETop and GIMME in the yeast batch fermentation model confirmed these differences. The CSN generated by GeNETop exhibited sufficient flexibility to accommodate metabolic transitions across the fermentation process. In contrast, the GIMME-derived CSN failed to converge during early fermentation and in the late stationary phase, highlighting its dependence on transcriptomic sampling times. In addition, comparative flux analysis revealed differences in compartmentalized metabolic strategies: GeNETop favoured mitochondrial pathways, whereas GIMME relied more heavily on nuclear or vacuolar reactions, suggesting alternative metabolic adaptations inferred by the two reconstruction approaches.

A relevant property of GeNETop is its reduced sensitivity to threshold selection. The multi-criteria reaction selection procedure combines flux variability, network topology, and transcriptomic information, preventing any single criterion from dominating the reconstruction process. For instance, reactions with low IVI scores may still be retained if they display high transcript expression or significant flux variability, thereby preserving pathways that may be required during metabolic transitions. By default, GeNETop employs a conservative variability threshold (*θ*_*v*_ = 1×10^−10^) to minimize the risk of excluding biologically relevant reactions. Sensitivity analysis showed that moderate variations of this threshold have a limited impact on the final CSN while preserving convergence in the dynamic simulations.

Beyond threshold selection, the influence of transcriptomic sampling times was also evaluated. The results indicate that GeNETop can compensate for limited transcriptomic resolution by retaining reactions supported by FVA and IVI criteria, reducing its dependence on specific transcriptomic sampling points. In contrast, the lack of convergence observed in the GIMME reconstruction at early time points suggests a stronger dependence on transcriptomic sampling resolution and threshold selection.

A key factor contributing to the robustness of GeNETop is the IVI metric, which integrates multiple centrality measures to capture the structural importance of reactions within the metabolic network and complements the flux-based information obtained through FVA. By combining IVI with transcriptomic information, the method preserves reactions that are both topologically relevant and transcriptionally supported. This multi-layer integration enables a balanced reduction of non-essential reactions while maintaining biological plausibility and computational efficiency in dynamic metabolic modelling.

## Conclusions

This work introduces GeNETop, a methodology for deriving context-specific metabolic networks tailored to dynamic environments. By integrating flux variability analysis (FVA), network topology metrics through IVI scores, and transcriptomic information, GeNETop addresses the challenge of generating context-specific networks that remain consistent under time-varying metabolic conditions.

The ability of GeNETop to construct context-specific metabolic models compatible with dynamic simulations from limited transcriptomic data represents an important advance in metabolic network reconstruction. This capability is particularly relevant for applications in industrial biotechnology and systems biology, where understanding dynamic metabolic shifts is essential for analysing cellular adaptation and guiding metabolic engineering strategies.

Beyond the specific case study presented here, the GeNETop framework may also facilitate the development of context-specific metabolic models for other dynamic biological systems, such as microbial communities or time-dependent physiological processes.

## Supporting information

**Supplementary File**. Pdf file including additional methodological details on the GeNETop workflow and supporting results for both the steady-state and batch fermentation case studies. These include model calibration, sensitivity analyses, IVI-based network characterization, and extended comparisons with alternative reconstruction methods.

**Supplementary Tables S1**. Excel file comparing cGIMME and GeNETop context-specific networks in continuous fermentation.

**Supplementary Tables S2**. Excel file with information related to the *S. cerevisiae* batch fermentation case. CSN derivation using GeNETop, transcriptomic data, and comparison with GIMME and FASTCORE.

**Supplementary Tables S3**. Excel file with information related to the *S. cerevisiae* batch fermentation case. Dynamic flux balance analysis results: comparison between GeNETop and GIMME. Metabolomics data.

All this Supporting Information necessary to explore the details of GeNETop is available on https://doi.org/10.5281/zenodo.18443625

## Acknowledgments

We thank Bas Teusink for stimulating discussions and feedback, and Artai R. Moimenta for providing the scripts for dynamic model calibration. This work has received funding from the European Union’s Horizon 2020 research and innovation programme under the Marie Sklodowska-Curie grant agreement No 956126; MCIU/AEI/FEDER grant reference PID2021-126380OB-C31 and PID2021-126380OB-C32 and Xunta de Galicia grant IN607B 2023/04. D.T-J. acknowledges funding by an Axuda de Apoio áEtapa Predoutoral GAIN–IN606A-2021/037.

## Notes

### Competing Interest Statement

The authors have declared no competing interest.

